# Non-human primate LIBRA-Seq accelerates neutralizing antibody discovery in RM vaccinated against HIV-1

**DOI:** 10.64898/2025.12.19.695353

**Authors:** Christopher T. Edwards, Aaron D. Silva-Trenkle, Anusmita Sahoo, Kendra Cruickshank, Stacey A. Lapp, Nagarajan Raju, Thang Ton, Amanda Metz, Emily McGhee, Faith A. Mbadugha, Tysheena P. Charles, Ankur Saini, Kiran Gill, Kathryn L. Pellegrini, Rui Kong, Jens Wrammert, Amit A. Upadhyay, Cynthia A. Derdeyn, Rama Rao Amara, Gabriel Kwong, Steven E. Bosinger

**Affiliations:** Division of Microbiology and Immunology, Emory National Primate Research Center, Emory University, Atlanta, GA 30329, USA; Wallace H. Coulter Department of Biomedical Engineering, Georgia Tech College of Engineering and Emory School of Medicine, Atlanta, GA 30332, USA; Department of Biological Sciences and Bioengineering, Indian Institute of Technology, Kanpur, Uttar Pradesh 208016, India; Department of Laboratory Medicine and Pathology, University of Washington, Seattle, WA, 98195, USA; Emory Vaccine Center, Emory School of Medicine, Emory University, Atlanta, GA, USA; Department of Pediatrics, Emory School of Medicine, Emory University, Atlanta, GA, 30329, USA; Emory National Primate Research Center Genomics Core, Emory National Primate Research Center, Emory University, Atlanta, GA 30329, USA; Department of Pathology and Laboratory Medicine, Emory School of Medicine, Emory University, Atlanta, GA, 30329, USA; Washington National Primate Research Center, University of Washington, Seattle, WA, 98195, USA; Department of Microbiology and Immunology, Emory School of Medicine, Emory University, Atlanta, GA, USA

## Abstract

Broadly neutralizing antibodies (bNAbs) exhibit protective efficacy against HIV-1 infection making them an ideal archetype for HIV-1 vaccine design. Presently, no vaccine candidate has induced antibody responses capable of meaningful protection against the swathe of circulating, difficult to neutralize tier 2 HIV-1 viruses. However, the development of stabilized, native-like envelope (Env) trimers such as BG505.SOSIP.664.T332N (BG505 SOSIP) has marked a significant advancement in vaccine design, due to their ability to elicit NAbs that neutralize tier 2 viruses in rhesus macaques (RM). NAb development following envelope trimer immunization in RM remains poorly understood, with hypothesized contributions from genetic variation at the IG loci, naive B cell repertoire, and differential gene expression in B cell lineages. To address these knowledge gaps, we have developed a set of BG505 SOSIP probes capable of recovering paired clonotype identity, antigen specificity, and gene expression of B cells in a high throughput fashion. These probes were constructed by conjugating biotinylated BG505 SOSIP to streptavidin covalently linked to both sc-RNA-Seq compatible DNA oligonucleotides and flow cytometry compatible fluorophores. Using these reagents, we isolated and sequenced BG505 SOSIP specific memory B cells from the PBMCs of an RM that developed high titers of neutralizing antibodies. To benchmark the accuracy of our technology, we compared our recovered heavy and light chain sequences to those identified from the same animal using conventional methodology and recovered 100% of previously identified NAbs. We then applied this technology to recover BG505 SOSIP specific memory B cells from five additional vaccinated RMs, cloned 34 antibodies for functional characterization, and identified ten antibodies with autologous neutralizing activity.

**Author Summary:** Understanding how effective antibodies arise after HIV vaccination is essential for developing a protective vaccine, yet studying these responses in non-human primates has been limited by low-throughput methods. In this study, we adapted a high-throughput single-cell sequencing approach to identify HIV envelope–specific antibodies from vaccinated rhesus macaques. This method allowed us to recover paired antibody sequences together with their antigen specificity from thousands of individual B cells. We successfully identified known neutralizing antibodies and discovered additional antibodies capable of neutralizing HIV across multiple animals. Our analysis revealed that vaccine-elicited antibody responses were dominated by a small number of expanded lineages, with shared genetic features among animals with stronger neutralization. These findings demonstrate that this approach can efficiently define the antibody repertoires generated by HIV vaccines and provide a powerful tool for evaluating and improving immunogens in preclinical vaccine studies.

## INTRODUCTION

In 2023, approximately 1.3 million people were infected with HIV-1, and over 600,000 people died from AIDS related illnesses(1). With millions of people living with HIV-1 across the world unable to access antiretroviral therapy and millions of others unaware of their HIV-1 status, developing a protective HIV-1 vaccine remains a central priority in the fight against the HIV-1 pandemic. However, the vast range of genetic variation in circulating strains, rapid establishment of long-lived latent reservoirs, and the prominent glycan shield that protects key neutralizing epitopes have proven to be formidable hurdles in the pursuit of an efficacious, antibody-based vaccine(2, 3). This is most clearly highlighted by the modest 31.2% vaccine efficacy of the only successful HIV-1 vaccine trial to date that was unable to be replicated in subsequent trials(4).

The failure of early vaccine trials to elicit neutralizing antibody responses prompted researchers to pivot to a “reverse vaccinology” approach(5–9). This strategy, broadly, is based on identifying monoclonal antibodies (mAbs) isolated from people living with HIV-1 with the ability to inhibit infection against a range of neutralization resistant (Tier 2) HIV-1 strains, and studying their biology (epitope specificity, germline allele, sequential accumulation of mutations) in order to inform the design of vaccine strategies capable of eliciting antibodies with similar properties. These antibodies known as “broadly neutralizing antibodies,” or “bNAbs,” are considered a critical correlate of protection against HIV-1 challenge, as they have been shown to both prevent simian – human immunodeficiency virus (SHIV) infection in rhesus macaques (RM) in passive antibody transfer studies and help maintain the suppression of HIV-1 during chronic infection humans(10–18). bNAbs arise in ∼10-30% of people naturally infected with HIV-1 (19–21), however eliciting them by vaccination has proven challenging and is one of the foremost priorities of HIV vaccine development(19–21). A vital advancement for the study of bNAbs was the development of thermostable, soluble HIV-1 Env trimers such as BG505.SOSIP.644.T332N (BG505 SOSIP)(22, 23). In pre-clinical immunization studies in macaques, BG505 SOSIP was able to elicit autologous NAbs against a tier 2 virus(24, 25). The BG505 SOSIP platform has advanced to clinical trials(26), and shown promise in eliciting B cell precursors to the VRC01 bNAb in humans(27). This construct has been used as an effective analytical tool to preferentially capture B cells producing bNAbs and elucidate key bNAb epitopes on Env(23, 24, 28). Our group has previously shown that RM immunized with BG505 SOSIP with 3M-052 adjuvant can confer protection against intravaginal challenge with BG505 SHIV(25). Though all vaccinated RM developed high levels of BG505 SOSIP binding antibody titers, only one third developed protective NAb titers. Analysis of the NAb responses from RUp16, an animal that developed inordinately high NAb titer, revealed the C3/465 glycan hole cluster as the immunodominant epitope among potent NAbs(29). Additionally, the serum from all but one of the animals protected from infection showed decreased neutralization capacity against a mutant BG505 SHIV-1 with the 465-glycan hole ocluded(29).

While RM have proven to be a highly valuable model for testing HIV-1 Env based vaccine constructs, conventional antibody sequencing techniques that rely on plate based single-cell sorting severely limit the throughput of NAb discovery. The recent development of LIBRA-Seq (linking B cell receptor to antigen specificity through sequencing) has been used to dissect humoral immune responses to pathogens and vaccine immunogens at the single cell level via barcoded antigens(30–42). In this study, we adapt the LIBRA-Seq platform to isolate BG505 SOSIP specific B cells from vaccinated RM *en masse,* identify public clones, inform the selection of 34 candidate mAbs for functional characterization, and ultimately identify autologous neutralizing antibodies.

## RESULTS

### Probe Design and Construction

To create a flow cytometry and LIBRA-seq compatible cell staining technology (**Fig 1A**), we conjugated streptavidin to Alexa fluorophores (AF) and sc-RNA-Seq compatible DNA oligonucleotides, which could be tetramerized with biotinylated proteins for staining of cells. First, we conjugated recombinant streptavidin with C-terminal cysteine to AF-maleimide at a 1 to 10 ratio. After removal of excess AF-meleimide through size exclusion spin filtration, streptavidin-AF were conjugated to DNA oligonucleotides through hydrazone chemistry and purified using size exclusion chromatography (**Fig. 1B**). Size exclusion chromatography results in distinct absorbance profiles between free streptavidin-AF647, free Oligo-1, and the streptavidin-AF647-Oligo-1 conjugate to allow for purification of conjugate with 7 to 12 ml of elution volume. We validated that the five different constructs constructed (AF647-Oligo-1, AF488-Oligo-2, AF647-Oligo-3, AF488-Oligo-4, and AF546-Oligo-5) have DNA conjugated to streptavidin monomers by protein gel, resulting in an additional band around the molecular weight of streptavidin monomer conjugated to DNA strand in both Coomassie and SYBR DNA gel staining (**Fig. 1C and Suppl. Fig. S1B**). To test the conjugation of the fluorophore, we used a murine tetramer system we had established previously(43), in which we had prepared streptavidin based Gp100-D^b^ tetramers. Here we created Gp100-D^b^ tetramers with streptavidin-AF or streptavidin-AF-DNA constructs to stain P14 splenocytes, resulting in similar staining profiles between DNA free and DNA conjugated streptavidin (**Fig. 1D and Suppl. Fig. S1A**). Our streptavidin-AF-DNA conjugates stained P14 splenocytes with similar efficiency to commercially available streptavidin.

**Figure 1.**
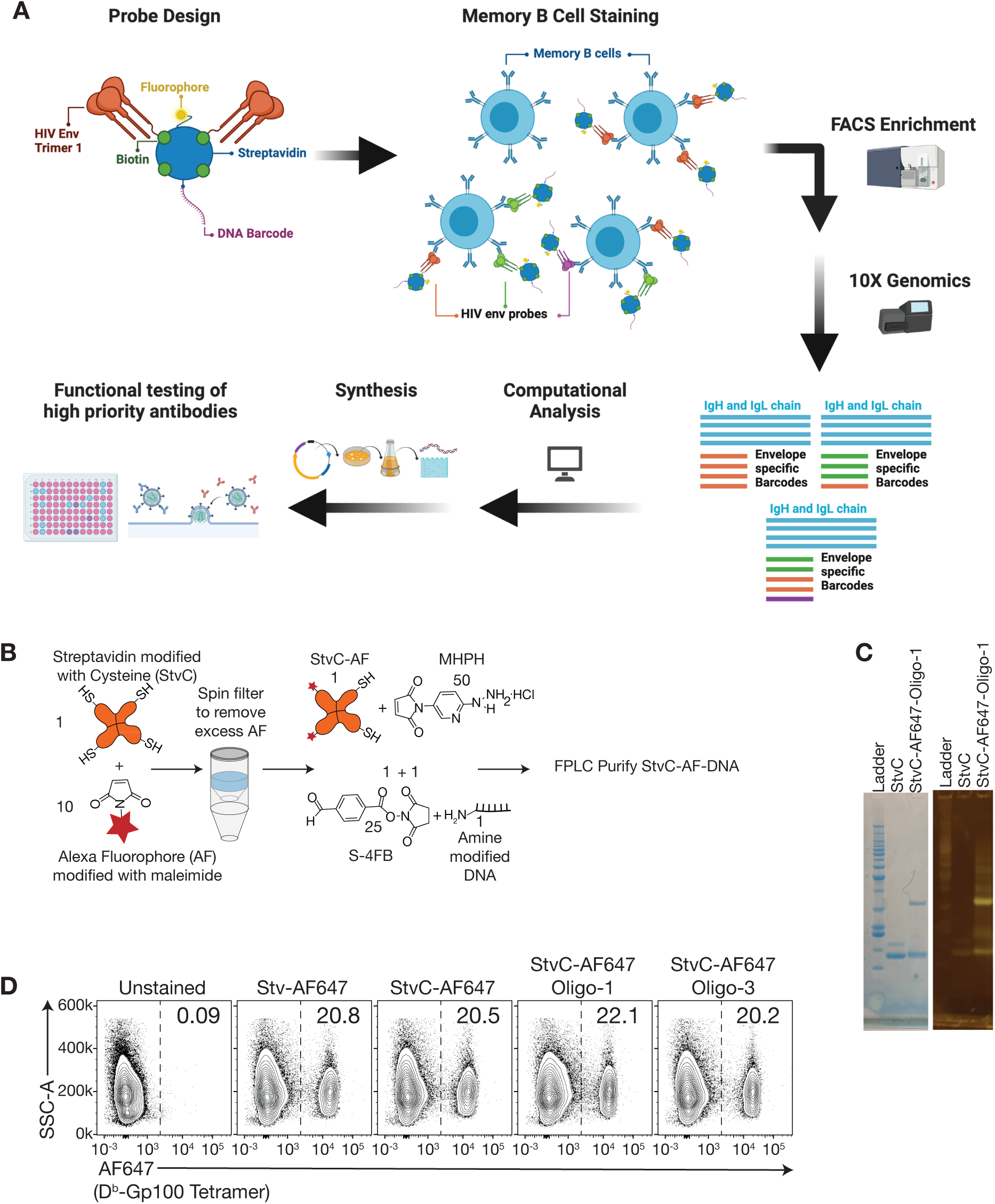
Schematic of the LIBRA-seq assay and probe design. (**A**) Schematic of the LIBRA-seq approach. Fluorophore and DNA oligo conjugated streptavidin are bound to biotinylated HIV-1 Env to create LIBRA-Seq probes. HIV-1 Env specific memory B cells bound to these probes are enriched via FACS prior to 10x capture. DNA libraries are generated from captured RNA and antigen barcodes. Bioinformatic analysis reveals binding profiles of individual memory B cells based on associated antigen barcodes and VDJ sequences. These profiles are used to prioritize clones for downstream functional characterization. (Created with <u>Biorender.com</u>) (**B**) StvC-AF-Oligo conjugates stain similarly to control Stv-AF. Streptavidin with N-terminal cysteine (StvC) was conjugated to maleimide modified alexa fluorophore (AF) at 10:1 ratio and excess removed by spin column purification before conjugating to amine modified DNA by hydrazone chemistry. StvC-AF-Oligo conjugates were purified from free StvC-AF and DNA by size exclusion chromatography. (**C**) Gel electrophoresis of StvC-AF-Oligo conjugates stained with Coomassie Blue (left) and SYBR DNA Gold (right). (**D**) Staining pmel splenocytes for Gp100-specific T cells with Db-Gp100-tetramers made from various streptavidin conjugates.

### NHP LIBRA-seq In Vitro Validation

To validate the BCR specificity and 10x single cell RNA-Seq compatibility of our probes, we utilized an engineered Ramos B cell line expressing VRC01, a CD4-binding-site-directed HIV-1 bNAb capable of binding BG505 SOSIP(30, 44, 45) (**Fig. 2A**). We mixed VRC01 B cells with parental RA.1 Ramos B cells that do not express VRC01 at 1:1, 1:100, and 1:1000 VRC01:RA.1 ratios and incubated with the BG505 SOSIP probe (**Fig. 2B**). Flow cytometry demonstrated that VRC01 expressing cells were recovered at the expected ratios, indicating highly efficient detection of VRC01. In addition to accurate detection of antigen specific VRC01 B cells via flow, we also performed independent 10x captures of RA.1 (9662 cells) and VRC01 (9128 cells) cells stained with the BG505 SOSIP probe. (**Suppl. Fig. S2C, Data File S1)**. Following QC filtering, 96.6% percent of VRC01 Ramos cells were found to express the canonical VRC01 CDRH3, while 94.1% of RA.1 Ramos cells expressed the parental RA.1 CDRH3 (**Suppl. Figs. S3A and 3B**). Despite the low frequency of BG505 SOSIP probe-positive RA.1 Ramos cells detected by flow cytometry, we observed a higher level of background antigen barcode reads in these cells following 10x capture (**Suppl. Fig. S2C, Data File S1)** compared to cells isolated via FACS prior to 10x capture (**Fig. 3G**).

**Figure 2.**
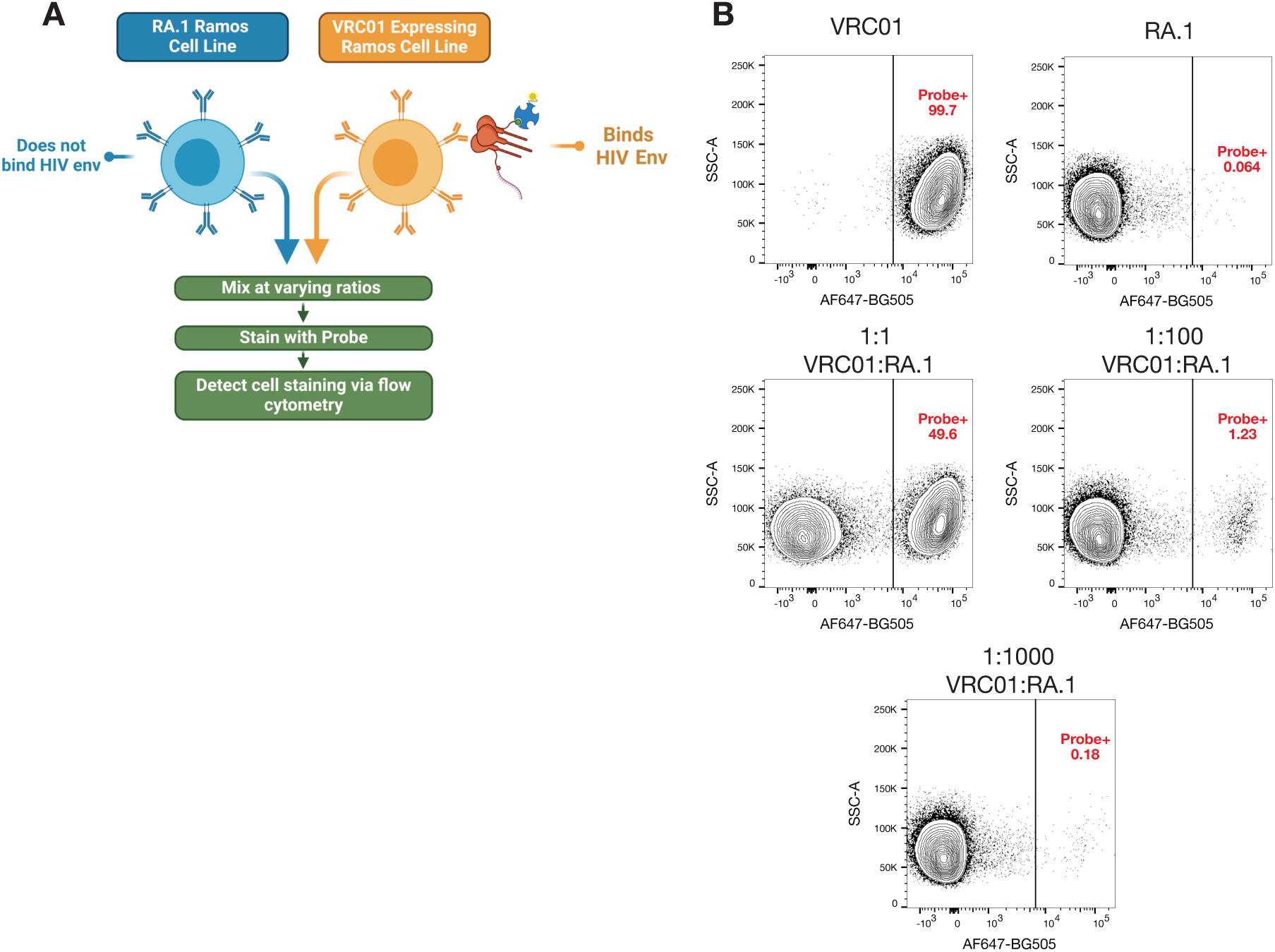
Validation of LIBRA-Seq compatible BG505 SOSIP probes *in vitro*. (**A**) Schematic of BG505 SOSIP based LIBRA-seq probe validation with bNAb expressing B cell lines. (Created with Biorender.com) (**B**) Binding of VRC01 or RA.1 expressing Ramos B cells to dual DNA-barcoded, fluorescently labeled BG505 SOSIP via flow cytometry. Surface bound VRC01 heavy chain expressing Ramos cells were stained either alone (top left) or at 1:1 (left middle), 1:100 (right middle), or 1:1000 (bottom) ratios with RA.1 expressing B cells. RA.1 Ramos cells were also stained alone (top right) to assess nonspecific binding.

**Figure 3.**
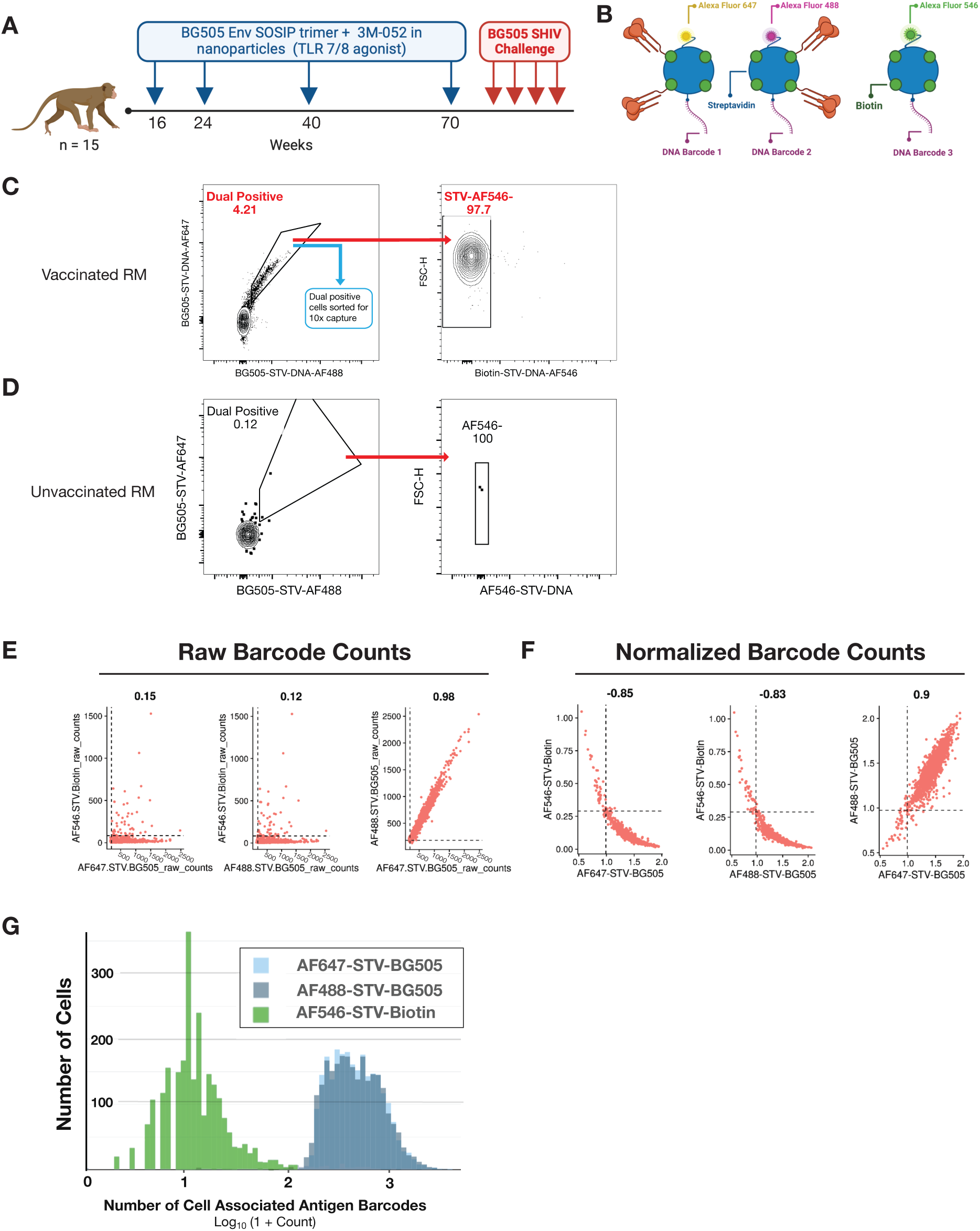
Benchmarking of NHP LIBRA-Seq compatible BG505 SOSIP probes *in vivo.* (**A**) Schematic representation of the immunization regimen. (Created with Biorender.com) (**B**) Design of LIBRA-Seq and flow cytometry compatible, BG505 SOSIP probes for dual antigen staining. Biotinylated BG505 SOSIP was conjugated to barcoded streptavidin linked to either AF647 (left) or AF488 (middle) fluorophores. Barcoded streptavidin bound to biotin only was linked to AF546 (right). (Created with Biorender.com) (**C**) Flow cytometry plots showing antigen specific memory B cells from cryopreserved PBMCs from RUp16 collected at weeks 87 and 90. Cells were gated on FSC and SSC characteristic of lymphocytes, singlets, live cells, CD3-, CD14-, CD16-, CD20+, CD27+, IgM-, IgG+, BG505-AF647+ and BG505-AF488+. (**D**) Representative flow cytometry plots showing BG505 SOSIP-specific memory B cells in an unvaccinated RM using the same gating strategy. Feature scatter plots highlighting the raw (**E**) and normalized (**F**) read counts for LIBRASeq barcodes, and show each combination of the biotin control and two BG505 SOSIP baits. The dotted lines represent the thresholds for antigen barcodes that were chosen empirically - 97^th^ percentile for biotin negative control and 3rd percentile for the two BG505 SOSIP barcodes. Each dot represents a unique cell. (**G**) Histogram displaying the number of cell associated antigen barcodes per LIBRA-Seq recovered B cell. Barcodes associated with BG505-AF647 are shown in blue, BG505-AF488 in grey, and Biotin-AF546 in green.

### NHP LIBRA-seq Recovers Known BG505 Neutralizing Antibodies from Vaccinated NHPs

We next sought to apply our LIBRA-Seq reagents to *ex-vivo* NHP samples. We have previously shown that immunization with BG505 SOSIP in RM provided significant protection against ten intra-vaginal challenges with BG505 SHIV-1(25). In this previous preclinical efficacy study, two groups of 15 RM received four subcutaneous immunizations with BG505 SOSIP in 3M-052 adjuvant, with one of these groups also receiving SIVmac239 Gag-expressing HVV to boost T cell responses, while a third group of 15 unimmunized RM received only 3M-052 adjuvant (**Fig. 3A**). Significant protection was observed in the SOSIP only and HVV + SOSIP vaccination groups compared to controls(25).

We used NHP LIBRA-Seq to identify the BG505 SOSIP specific antibody repertoire of RUp16, a RM that was protected from BG505 SHIV-1 challenge and developed the highest NAb titer (ID50 = 6068). This animal was selected for LIBRA-Seq benchmarking as it had undergone previous high-resolution analysis of NAb associated with high titer and protection using conventional methodology(25, 29). PBMCs isolated from RUp16 at weeks 89 and 92 were combined and stained with dual BG505 SOSIP probes and a negative bait probe with unique fluorophores and corresponding DNA oligos (**Fig. 3B and 3C**). To assess congruency between FACS and 10x readouts, we chose to sort dual positive memory B cells regardless of negative bait binding. Over 13,000 dual BG505 SOSIP probe bound memory B cells (defined as FSC and SSC characteristic of lymphocytes, singlets, live cells, CD3-, CD14-, CD16-, CD20+, CD27+, IgM-, IgG+, BG505-AF647+ and BG505-AF488+) were sorted for 10x capture, representing approximately 4.21% of the circulating IgG+ memory B cell population. Of the dual-positive cells sorted for 10x capture, 97.7% were found to be negative for the biotin bound LIBRA-Seq probe by FACS (**Fig. 3C**). In contrast, only 0.12% of memory B cells isolated from an unvaccinated RM were found to bind to BG505 SOSIP probes (**Fig. 3D**). Following 10x capture and subsequent library generation of RUp16’s antigen specific memory B cells, we were able to recover both the BCR sequence and antigen barcode libraries from 1706 cells, with 1643 (96.3%) associated with both BG505 SOSIP barcodes and lacking any biotin bait barcode (**Table 1**). These data highlight the consistency between our FACS generated antigen binding profiles and those generated from the 10x digital readout of antigen associated barcodes.

**Table 1.** Sample Details. PBMCs were isolated from whole blood of BG505.SOSIP vaccinated rhesus macaques at various timepoints following 2-4 immunizations. PBMC samples were pooled from two timepoints prior to memory B cell enrichment.

| Animal Code | Weeks after initial immunization WB was sampled | Total Number of PBMCs (millions) | ID50 Titer (week 26) | # of Sorted B cells for 10x Capture | 10x Recovered Cells | Cells with Antigen Barcodes and VDJ Recovered | Cells Positive for both BG505 Probe Barcodes and Negative for Biotin Barcode |
| --- | --- | --- | --- | --- | --- | --- | --- |
| **275_12** | 36+43 | 19.5 | 334 | 180 | 52 (28.89%) | 41 | 36 |
| **RYs15** | 40+43 | 14.5 | 41 | 634 | 62 (9.77%) | 29 | 27 |
| **RHe16** | 36+43 | 22.5 | 187 | 640 | 94 (14.69%) | 45 | 41 |
| **Rlr15** | 36+43 | 28 | 529 | 2089 | 428 (20.49%) | 223 | 216 |
| **RLk15** | 36+43 | 18.5 | 501 | 280 | 86 (30.71%) | 63 | 61 |
| **RUp16** | 89+92 | 13.3 | 6068* | 13069 | 2792 (21.36%) | 1706 | 1643 |
| *week 82 titer |  |  |  |  |  |  |  |

For each memory B cell with successfully recovered BCR and antigen barcode libraries, the LIBRA-seq scores for each BG505 SOSIP probe and negative bait probe were calculated based on the number of unique molecular identifiers (UMIs) detected for each construct (**Fig. 3E-3F**). Raw counts for each BG505 SOSIP probe were highly correlated with one another (Pearson’s r = .98) (**Figs. 3E-3G, Data File S2**) and maintained a high correlation following normalization for total number of reads per cell (Pearson’s r = .9) (**Fig. 3F**). Using the original LIBRA-seq analysis pipeline, we found that many cells were assigned LIBRA-Seq scores skewed by the ratios of raw antigen barcode counts due to scaling influenced by cells with higher overall numbers of recovered barcodes. Our dual positive and single negative probe schema necessitated additional oversight when determining antigen specificity beyond LIBRA-Seq score cutoffs. Thresholds for antigen specificity based on antigen barcode read counts were chosen empirically - 97th percentile for biotin and 3rd percentile for the two BG505 SOSIP barcodes. Using these thresholds, cells were classified as positive if the normalized values were surpassed these thresholds for both BG505 SOSIP antigens and but not for biotin.

To further benchmark our NHP LIBRA-Seq approach, we compared the BCR sequences of the recovered BG505 SOSIP specific memory B cells to the published heavy and light chain sequences of BG505 SOSIP specific mAbs isolated from RUp16(29). We recovered 24 of the 48 previously published heavy chains and 40 of the 44 previously published light chains of BG505 SOSIP binding mAbs within 85% sequence identity (**Fig 4A, Suppl. Fig. S6B**). Among the overlapping sequences were the heavy and light chains of all four BG505 SOSIP neutralizing mAbs previously isolated from RUp16 with BG505 gp120 monomers(46). Clonal analysis revealed 302 shared clonotypes of LIBRA-Seq recovered heavy chains, including 22 out of 48 previously described heavy chains, with all four mAb heavy chains represented in the second largest clonal family (**Fig 4B and 4C**). The results from vaccinated RM RUp16 suggest that the LIBRA-seq platform can be successfully applied to the NHP model to identify antigen specific mAb characterized using conventional cloning methodology.

**Figure 4.**
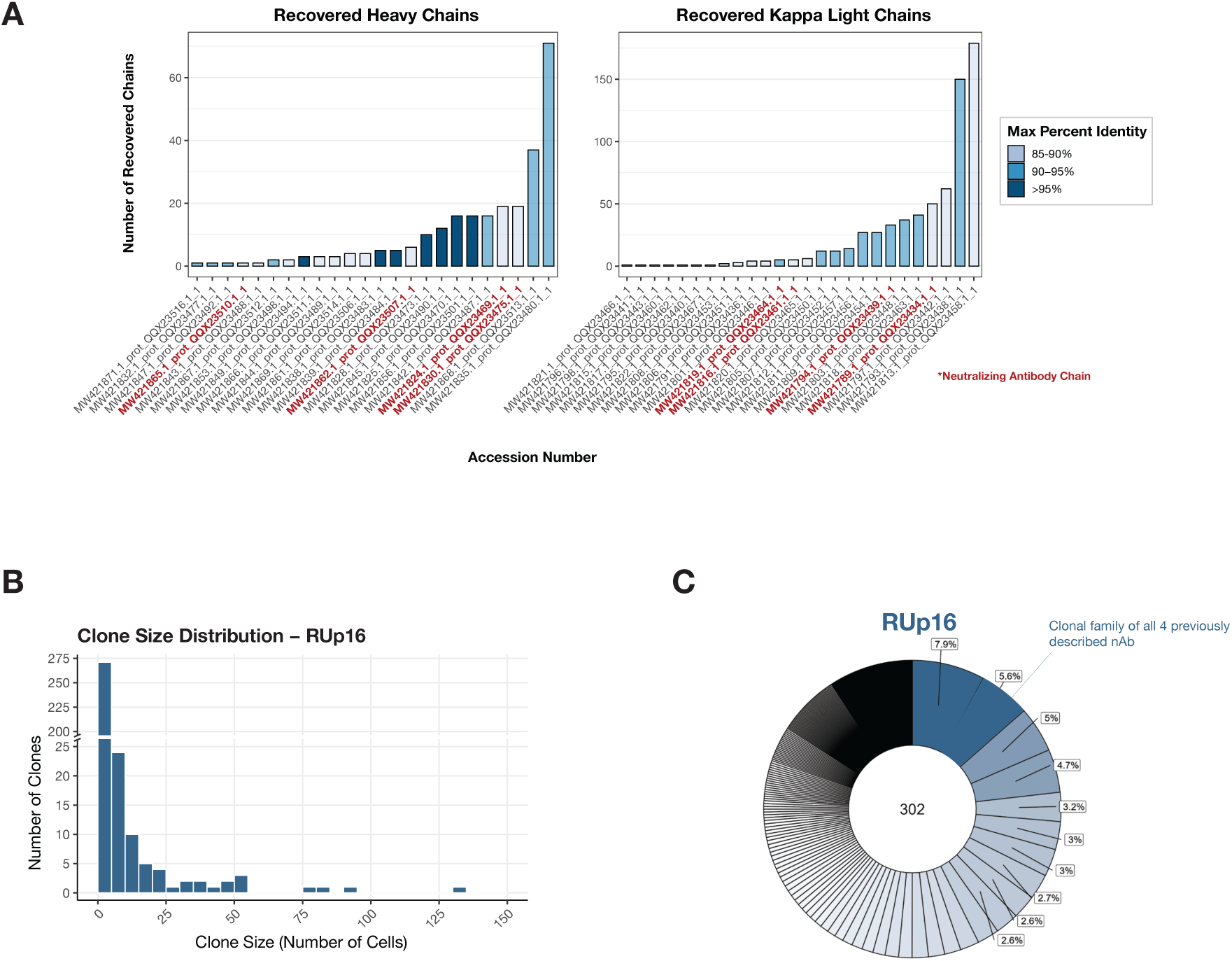
LIBRA-Seq identifies BG505 Env neutralizing antibody lineages from animal RUp16. (**A**) Heavy chains (left) and kappa light chains (right) recovered from RUp16 using LIBRA-Seq that map to previously identified BG505 binding light chain sequences. Shared chains were determined by same V and J gene usage and CDR3 sequence identity above 85%. Color denotes the maximum percent sequence identity shared. Heavy and light chains matching the four neutralizing monoclonal antibodies identified previously by Charles et al. are displayed in red (29). Distribution of clone sizes (**B**) and proportion (**C**) of LIBRA-Seq recovered BG505 SOSIP specific B cells from RUp16. The number in the center (**C**) reflects the total number of B cell clonotypes identified as BG505 SOSIP specific with LIBRA-Seq with clones ranked clockwise from the top center in order of relative frequency. The frequency of the top 10 most abundant clones are indicated, with the most frequent clones noted in dark blue. B cells were considered clones through shared V-genes, J-genes, identical CDR3 length and greater than 70% CDR3 nucleotide sequence identity for both heavy and light chains.

### NHP LIBRA-seq Accelerates Discovery of Neutralizing mAbs from BG505 Vaccinated RM

We next sought to apply LIBRA-Seq to 5 additional RM vaccinated in the same study as RUp16. Animals were selected based on their protection from NAb challenge and had ID50 titers ranging from 41 to 529 (**Table 1).** Using the same memory B cell panel and LIBRA-Seq probes as described above, BG505 SOSIP specific memory B cells were sorted from weeks 36 to 43, representing the circulating memory B cell population prior to and following the third boost with BG505 SOSIP. Additionally, only memory B cells negative for the biotin bound bait were sorted for 10x capture from these RM (**Suppl. Fig. S4A, Data File S2**). Antigen specific cells represented 0.34% to 0.8% of the isolated memory B cells in these RM, lower in comparison to RUp16’s 4.21%, likely because these cells were isolated prior to the final boost, as well as the observed differences in ID50. Despite the lower frequencies of antigen specific cells in these samples, we recovered the BCR sequences and associated antigen barcodes for a total of 401 memory B cells from these 5 RM (**Table 1**).

To confirm LIBRA-Seq’s accuracy in identifying antigen-specific B cells, we produced five antibodies with high BG505 SOSIP associated LIBRA-Seq scores per animal (n=30), and an additional four antibodies with high biotin bait associated scores to assess LIBRA-Seq’s ability to flag non-specific memory B cells. Antigen specificity as predicted by LIBRA-seq was validated by ELISA. mAbs PGT151 and PGT145, that bind to trimeric epitopes in BG505 SOSIP were run as positive controls and influenza HA specific mAb EM4C04 was run as a negative control (**Fig. 5A**). All 30 antibodies with LIBRA-Seq scores denoting a high specificity for BG505 SOSIP exhibited binding via ELISA, with LIBRA-Seq scores trended with ELISA area under the curve (AUC) values (**Fig. 5B-5C**). Interestingly, of the four antibodies with high LIBRA-Seq scores for the negative bait probe, two were observed to bind BG505 SOSIP, while the remaining two exhibited AUC values below the limit of detection. One of these two antibodies, Rup16_158_TriplePostive, had a related clone amongst the dual positive set, RUp16_158_DualPositive, that also demonstrated strong BG505 SOSIP binding via ELISA. These two antibodies were found in a total of 9 memory B cells of the same clonotype, and the Rup16_158_TriplePostive sequence originated from the only memory B cell with significant negative bait barcodes within that clone.

**Figure 5.**
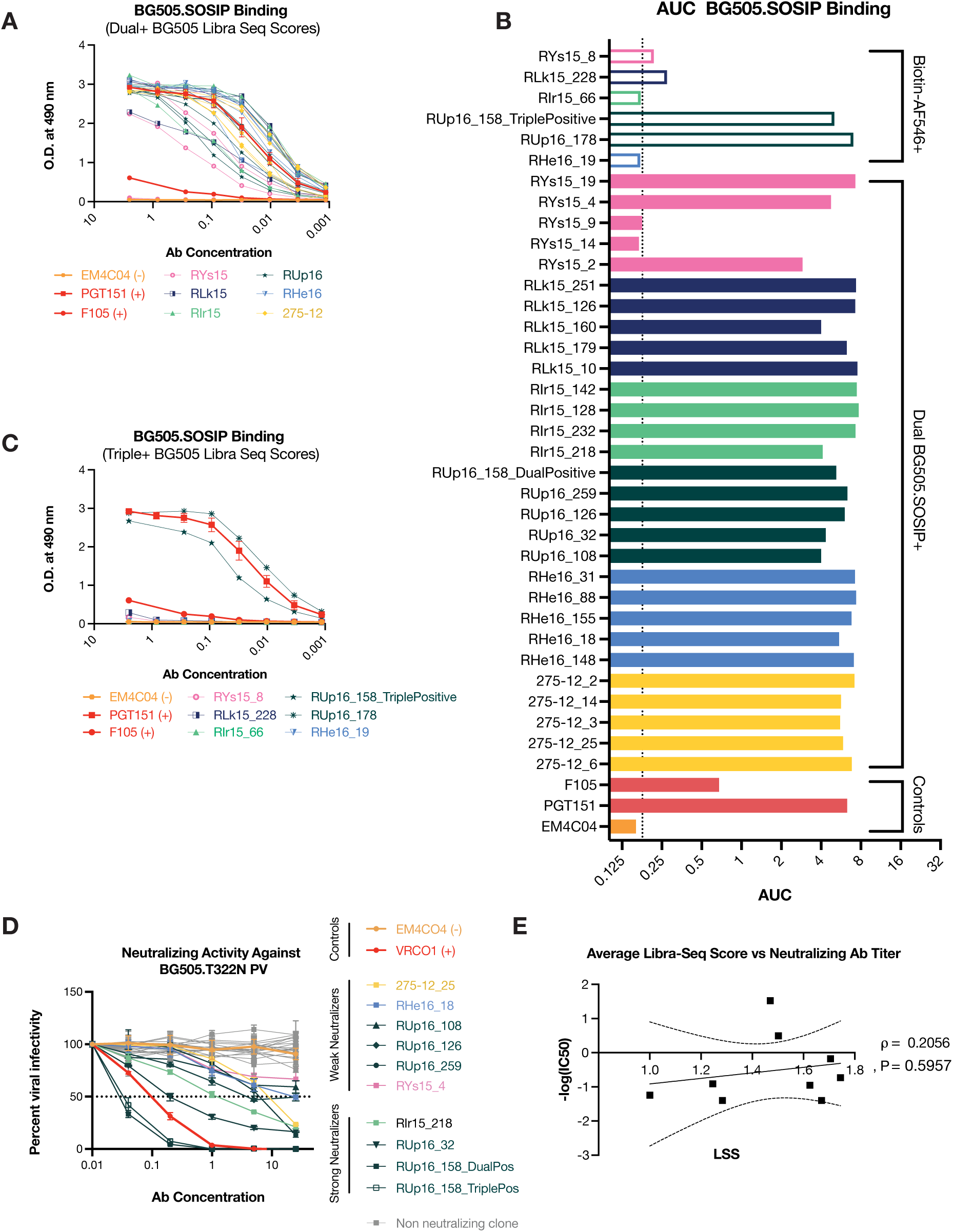
LIBRA-Seq accurately identifies BG505 SOSIP binding mAbs. Quantitative analysis of binding for affinity to BG505 was evaluated and depicted in a binding curve (**A** and **C**) or bar graph (**B**). A three-fold dilution of each mAb was done, starting at 5 ug/ml plotted based on the OD at 490nm. Bars represent the area under the curve for each mAb (x axis). The dotted line marks the three times the background signal of samples dilution buffer. Previously characterized HIV-1 specific human IgG1 antibodies (F105 and PGT151) were used as a positive control. An influenza HA-specific human IgG1 antibody (EM4C04) was used as a negative control. Binding affinities were calculated for both dual BG505 probe positive mAbs (**A**), and triple positive, BG505-SOSIP and biotin probe binding cells (**C**). (**D**) Quantitative analysis of neutralizing activity against BG505.N332 pseudoviruses in the TZM-bl assay was evaluated and depicted in infectivity curves. The percent virus infectivity is shown on the y axis plotted against the monoclonal antibody concentration on the x axis on a log10 scale. Serially diluted antibody at 25, 5, 1, 0.2, and 0.04 μg/mL was tested. (**E**) Correlation of average BG505-SOSIP LIBRA-Seq score with BG505.N332 pseudovirus neutralization titer. Dotted lines depict 95% confidence intervals.

We then sought to assess the neutralizing capacity of the recovered antigen specific antibodies. Of the 32 antibodies confirmed to bind BG505.SOSIP via ELISA, 10 isolated across 5 animals were found to exhibit neutralizing activity against BG505.N332 pseudovirus (**Fig. 5D**), with IC50 values ranging from 0.03 ug/ml to >25ug/ml. Of these 10, four demonstrated notable IC50 values (IC50<= 1.5 ug/ml), with the three strongest neutralizers all originating from RUp16. The IC50 curves of both RUp16_158 clones were highly similar, demonstrating strong neutralizing activity surpassing that of positive control bNAb, VRC01.

### NHP LIBRA-seq Analysis of BG505.SOSIP Vaccination Reveals a Highly Oligoclonal Repertoire with Shared Lineages

We then investigated the properties of the antigen specific memory B cell repertoires recovered with NHP LIBRA-Seq. Clonal analysis revealed an oligoclonal repertoire, with higher clonality correlating with higher serum ID50 titers (**Fig. 6A and 6D, Data File S3**). Across the cohort, the top 10 clones represented 31% to 50.3% of the of the total antigen specific memory B cell pool. We observed public heavy chain V gene usage across the cohort, with IGHV4-79 representing the most highly shared allele amongst BG505 SOSIP specific memory B cells (**Fig. 6B**). Interestingly, the NAb lineage contributing most significantly to RUp16’s high neutralizing titer also utilizes IGHV4-79 in its heavy chain. We also observed 6 public clones shared by pairs of vaccinated RM (**Fig. 6C**). Frequency of somatic hypermutation (SHM) was analyzed at both the per-cell and per-clone levels for IgH, IgK, and IgL to assess the extent of mutation in memory B cells recovered from BG505 SOSIP-vaccinated RM (**Fig. 7A and 7B**). The analysis revealed variation in SHM frequencies across animals, with the highest mutation rates observed in RUp16, consistent with both its sampling at a later timepoint and high NAb titers. SHM was observed at a frequency of 0.045-0.066 in IgH, 0.022-0.039 in IgK, and (0.025 - 0.044). These findings highlight the diverse characteristics of the antigen-specific memory B cell repertoires revealed through LIBRA-Seq and their potential relationship to the generation of potent neutralizing antibody responses in vaccinated RM.

**Figure 6.**
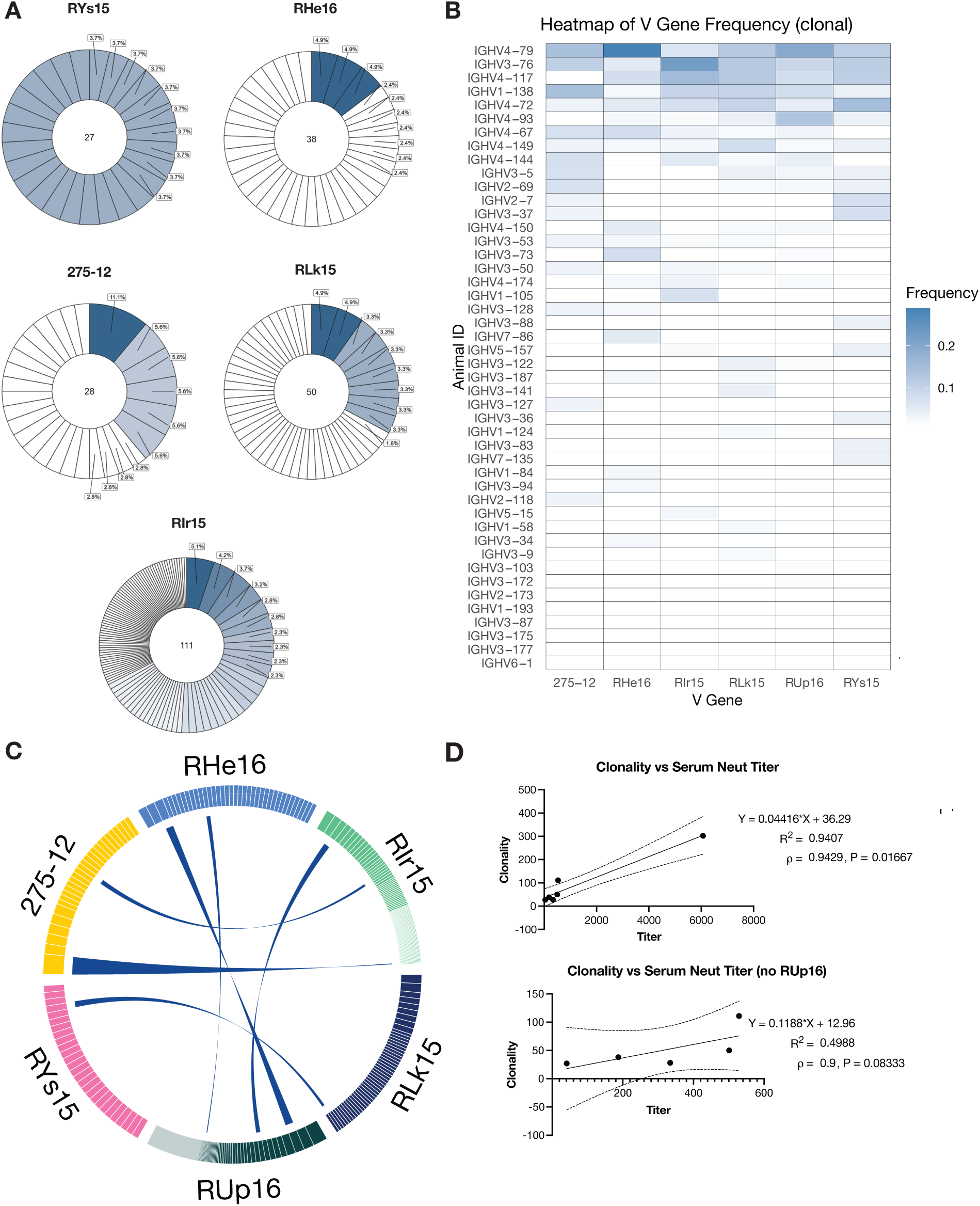
LIBRA-Seq identifies shared clonotypes of BG505 SOSIP specific B cells from vaccinated RM. (**A**) Clonal expansion of BG505 SOSIP specific B cells depicted in donut charts. Each donut reflects the total number of B cell clonotypes identified as BG505 SOSIP specific with LIBRA-Seq (center) from each RM and the relative frequency of each individual MBC clonotype. The frequency of the top 10 most abundant clones are indicated, with the most frequent clones noted in dark blue. (**B**) Heatmap of V gene frequency per RM, ranked from most shared V genes at the top to least shared at the bottom. Higher frequency is noted in dark blue. (**C**) Circos plot displaying shared BG505 SOSIP specific memory B cell clonotypes among six RM vaccinated with BG505 SOSIP. Each segment represents an individual animal, with connecting ribbons indicating shared clonotypes between animals. The thickness of each ribbon corresponds to the frequency of the clone. B cells were considered clones through shared V-genes, J-genes, identical CDR3 length and greater than 70% CDR3 nucleotide sequence identity for both heavy and light chains. Correlations of clonality with serum BG505 pseudo virus neutralization titer for all animals (top) and all animals without Rup16 (bottom). Dotted lines depict 95% confidence intervals.

**Figure 7.**
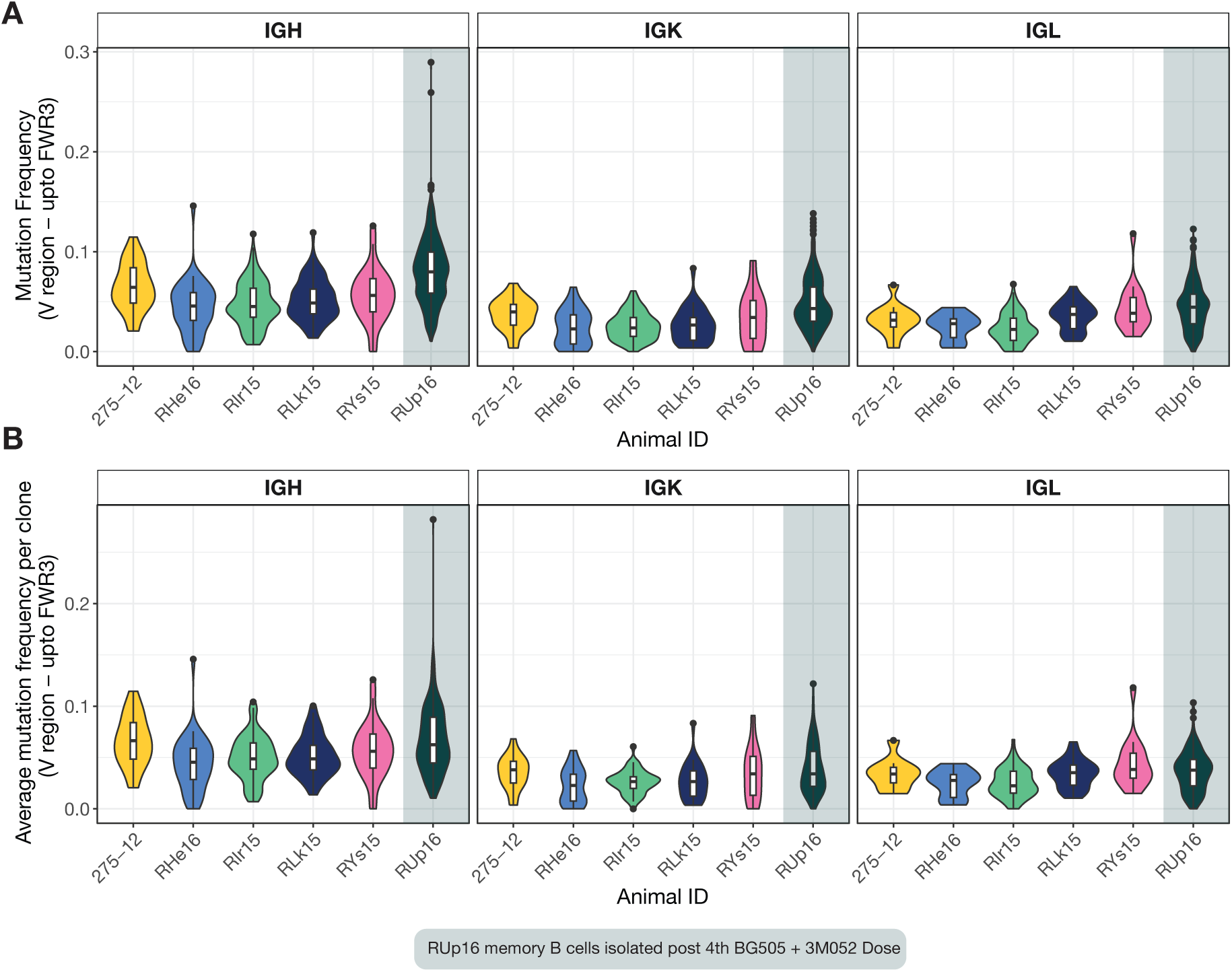
LIBRA-Seq recovers somatically hyper mutated BG505 SOSIP specific memory B cells. (**A**) Somatic hyper mutation (SHM) frequencies per LIBRA-Seq recovered BG505 SOSIP specific memory B cell across Ig heavy chain CDR3 (left), and Ig light chain IgK (middle) or IgL (right). Violin plots show the distribution of mutation frequencies for each animal, with overlaid boxplots indicating medians and interquartile ranges. (**B**) Mean mutational frequency per clone was calculated by averaging the mutation frequencies of all sequences assigned to the same clonal lineage. Violin plots display the distribution of average SHM frequencies per clone for each animal and locus with overlaid boxplots indicating medians and interquartile ranges. B cells were considered clones through shared V-genes, J-genes, identical CDR3 length and greater than 70% CDR3 nucleotide sequence identity for both heavy and light chains.

## DISCUSSION

To date, only one HIV-1 vaccine regimen has been shown to modestly protect humans from HIV-1 infection, an effect that has not been replicated elsewhere(3, 47). Recent innovations in immunogen design, delivery, and adjuvants have yielded breakthroughs in eliciting autologous, NAbs against tier 2 viruses and important bNAb precursors in humans, and protect against challenge with the matched strain in RM(24, 48–52). Here, we adapted the recently developed LIBRA-Seq platform to be compatible with the preclinical non-human primate model. We applied LIBRA-Seq to RM vaccine study samples as a proof of concept for studying memory B cell responses to immunogens in a high resolution, high throughput manner. We focused on RM vaccinated with BG505 SOSIP that were protected from infection with high serum NAb ID50 titers(25). Previous work had shown that clade A BG505 SOSIP immunogens elicit a range of neutralizing titers in the RM model, primarily targeting the C3/465 glycan hole cluster(29).

In this study, we were able to apply LIBRA-Seq and recapitulate many of the previous findings characterizing BG505 SOSIP vaccination of NHPs using conventional methodology. We were also able to expand on previous knowledge, recovering the sequences of BG505 SOSIP specific cells across multiple animals in a high throughput manner. In addition to recovering the previously identified neutralizing clones, we were able to identify shared gene usage and clonotypes across multiple animals. LIBRA-Seq scores also served as an additional metric for prioritizing clones for functional validation, with all selected clones exhibiting high BG505 SOSIP binding titers. From the 2107 cells successfully isolated with complete VDJ and surface barcode libraries across our cohort, we successfully recovered ten antibodies with autologous neutralizing activity, three exhibiting strong neutralizing activity. We attribute this high success rate in part to our selection being functionally informed by absolute barcode count and LIBRA-Seq score, which enabled the identification of clones with strong patterns for BG505 specificity across individual cells. Though our study only utilized 2 unique probes, we show how LIBRA-Seq can enhance the resolution of the analysis of the vaccine elicited B cell repertoire even with the limited modality. Future studies focusing on a panel of immunogens to probe epitope specificity are necessary to utilize the technology to its full potential.

Recent clinical studies have investigated immunogens specifically designed to elicit responses from bNAb precursors(53–55). Such studies represent the first step in guiding antibody maturation toward broadly neutralizing lineages, a process referred to as germline shepherding. In such studies, conventional analysis of binding and neutralizing titers lack insight into the gene sequences of elicited B cell lineages. LIBRA-Seq is uniquely poised to support and accelerate such studies by enabling high-throughput recovery of antibody specificity and gene usage, which is especially critical for analyzing and tracking precursor lineages over the course of vaccination, as well as identifying shared clonotypes across animals. To date, LIBRA-Seq has successfully recovered NAb from the convalescent plasma of a convalescent COVID-19 donor, subjects living with HIV-1, B cells elicited by the BNT162b2 vaccine in COVID-19 unexperienced and experienced individuals, and has also recovered public clonotypes in a guinea pig model of HIV-1 vaccination, but has yet to be applied to RM models of HIV-1 vaccination (30, 31, 35). The utility of LIBRA-Seq in identifying shared neutralizing epitopes across viral species has recently been shown in a preprint analyzing antibody responses elicited in the HIV Vaccine Network (HVTN) 124 study, a clinical trial using a polyvalent HIV-1 vaccine design, which isolated several glycan reactive and Fab-dimerized glycan-reactive antibodies capable of broad HIV-1 and even HCV neutralization. Such work illustrates necessity of high throughput approaches capable of resolving the epitope specificity of antibody lineages of interest across a vaccine cohort. With our data, these findings highlight how LIBRA-Seq can support next-generation, clinically implemented, germline-shepherding strategies by accelerating the identification of convergent antibody specificities, shared neutralizing epitopes, and public clones,

Overall, this study demonstrates the utility of LIBRA-Seq in enhancing the resolution of vaccine elicited B cell repertoire analysis in the RM model. By enabling high-throughput recovery of antigen-specific sequences, LIBRA-Seq provides a powerful approach for investigating the genetic determinants of antibody responses(30, 32–42). As the field advances toward precision immunogen design, applying LIBRA-Seq to larger cohorts and diverse immunogen panels will be essential for optimizing germline-targeting strategies and improving HIV-1 vaccine efficacy.

## EXPERIMENTAL MODEL AND SUBJECT DETAILS

### Ethics Statement

The RM immunization experiment from which the serum samples were derived have been described previously(25). The study was approved by the Institutional Animal Care and Use Committee (IACUC) at Emory University and followed NIH guidelines. Animal research was also in compliance with the Animal Welfare Act and other Federal statutes and regulations relating to experiments involving animals. All animal research adhered to the principles stated in the 2011 Guide for the Care and Use of Laboratory Animals prepared by the National Research Council. Emory National Primate Research Center (ENPRC) is fully accredited by the Association for Assessment and Accreditation of Laboratory Animal Care (AAALAC). Methods of euthanasia were consistent with the American Veterinary Medical Association with Guidelines.

### Cell Lines

Surface VRC01 expressing Ramos B cells were provided by Dr. Daniel Lingwood at the Ragon Institute of MGH, MIT and Harvard. This cell line was generated and cultured as previously described(45), and validated for binding to our antigen probes by FACS (Figure S1B). RA.1 ramos cells were obtained from ATCC and cultured according to manufacturer instructions.

### Animal Models

PBMC samples were obtained from a total of 6 Indian rhesus macaques (*Macaca mulatta*) immunized during a vaccine efficacy study carried out previously at the Yerkes National Primate Research Center(25). The study utilized female RM that were 3–15 years of age and confirmed negative for SIV infection. The immunization regimen has been previously described(25).

## MATERIALS AND METHODS

### Expression and purification of trimeric BG505.SOSIP.664 T332N-avi-biotinylated protein

The BG505 SOSIP Env insert (Genebank id ANG65466.1, res. 31-664, A501C/T605C/T332N, 508RRRRRR511) was synthesized by Genescript with (a) GMCSF leader sequence (MWLQGLLLLGTVACSIS) at its N-terminus end; (b) GTGS linker sequence and Avi tag (GLNDIFEAQKIEWHE) at its C-terminus. The insert was sub-cloned between ClaI and NheI sites of pGA1vector (KanR). NEB® 5-alpha E. coli cells (NEB, catalog no C2987H) and Sanger sequencing were used to transform, screen and confirm the positive clones respectively. The envelope protein cloned in pGA1 plasmid was expressed along with furin (expressed from an AmpR plasmid provided by Prof. John P. Moore), by transient transfection of Expi293F cells, in the ratio 4:1(22) using the ExpifectamineTM 293 transfection kit (ThermoScientific) as per manufacture’s protocol and grown at 37oC, 8% CO2 at 130rpm. The purification process used here has been previously described(56). Briefly, the supernatant was harvested 72hrs after transfection in presence of EDTA free protease inhibitor (Millipore Sigma, catalog no 11836170001) and affinity purified by lectin agarose (Vector Labs, catalog no AL-1243-5, pre-equilibrated with PBS). Bound protein was eluted in presence of 1M methyl a-D-mannopyranoside (Sigma). The protein was dialyzed against PBS and subjected to size-exclusion chromatography using a Superdex 200 Increase 10/300 GL (Sigma, GE Healthcare product) column on an AktaTM Pure (GE) system. The trimeric peak was collected, concentrated using Amicon Ultra-4, MWCO 100kDa, and quantified by BCA assay (PierceTM, ThermoScientific). The trimeric status and purity of the protein was confirmed by BN-PAGE (NuPAGETM,4-12%BisTris Protein Gels, ThermoScientific). The protein was concentrated to ∼9mg/ml. Reaction mixture containing BG505.SOSIP.664-avi protein, BirA (25µg for 10nmole of avi tagged protein), BiomixA (10X), BiomixB(10X) was incubated at 30oC for 45min, as per manufacture’s protocol (BirA500, Avidity). Free biotin was removed by passing the reaction mixture through a Amicon Ultra-4, MWCO either 100kDa. The protein was found to be ∼90% biotinylated, as estimated by ELISA using standards (MBP, MBP-avi-biotinylated) provided with the kit. The trimeric status of the biotinylated protein was confirmed by BN-PAGE. The protein was stored at 1mg/ml concentration.

### Memory B cell Immunophenotyping

(AMQAX1000). Appropriate volume for 1 million cells and 5 million cells were added to FACS tubes for control and flow panel respectively. Samples were centrifuged at 180g for 5 minutes then decanted. Control samples were immediately fixed and resuspended in 300µL of 1% PFA then stored in 4°C until ready for flow acquisition. Samples were incubated with 100µL of Fixable Viability Dye eFluor506 mix (65-0866-14) from eBioscience (1:1000 dilution) for 20 minutes at room temperature (RT) in the dark. Followed by a wash step with 2mLs of BSA Stain Buffer (554657), centrifuged at 300g for 5 minutes then decant. Samples were then incubated with 100µL of FC Block mix (14-9165-42) from Invitrogen (1:50 dilution) for 30 minutes at RT in the dark and followed by a wash step. Samples were then incubated with 100µL of biotin-quenched conjugated BG505 AF647/AF488 probe mix (1:1 dilution) for 30 minutes at RT in the dark and followed by a wash step. Samples were then incubated with 100µL of biotin-quenched unconjugated probe mix for 30 minutes at RT in the dark and followed by a wash step. Samples were then incubated with 100µL of the stain mix for 30 minutes at RT in the dark using the following mAbs: IgG BV650 (clone G18-145; 1.0µL; cat # 740596); IgM PerCP-Cy5.5 (clone G20-127; 3.0µL; cat # 561285); CD3 PE-CF594 (clone SP34-2; 2.0µL; cat # 562406); CD14 PE-CF594 (clone MφP9; 1.0µL; cat # 562334); CD16 PE-CF594 (clone 3G8; 1.0µL; cat # 562320) from BD Bioscience; CD27 BV421 (clone O323; 2.5µL; cat # 302824); CD20 APC-Cy7 (clone 2H7; 2.0µL; cat # 302314); BSA Stain Buffer (87.5µL; cat # 554657) from Biolegend; followed by a wash step. Samples were fixed and resuspended in 500µL of 1% PFA for cytometry acquisition. RAMOS cells were gated based on their FSC and SSC characteristics, singlets, live cells, CD3/CD14/CD16 (-) and CD20 (+), CD27 (+) and CD20 (+), IgM (-) and IgG (+). These cells were then assessed for their affinity to BG505 probes. Samples were run on BD FACSymphony A5 driven by FACS DiVa software and analyzed with FlowJo (Version 10.10).

The same protocol was performed on PBMCs from Rhesus Macaques with the exception of the centrifuge speed at 300g for 10 minutes. Additionally, PBMCs were resuspended and strained through 70µM cell strainers in pre-chilled R10 media for sorting on BD FACS Aria BD CompBeads Anti-Mouse Ig, κ/Negative Control Compensation Particles Set (cat # 552843) were used for single fluorophore stains to select for the brightest peak. Compensations were prepared fresh and acquired for each assay.

### Oligonucleotide barcodes

69 base pair (bp) in length oligonucleotide barcodes were designed with the following structure: 5’-5AmMC12-Read 2N-N10-Feature Barcode-N9-Capture Sequence-3’, where the 5AmMC12 represents a 5’ amino modification and 12 carbon linker for conjugation to streptavidin, the Read 2N is the Truseq read 2 sequence, 5’-CGGAGATGTGTATAAGAGACAG-3’, N10 and N9 denote random nucleotide sequences of 10 and 9 bp in length respectively and are used as universal molecular identifiers (UMIs), the Feature Barcode is a known 15 bp sequence selected from the 10x feature barcode whitelist, and the capture sequence, 5’-CCCATATAAGA*A*A-3’ is required for annealing to Chromium Next GEM Single Cell 5’ Gel Beads (V2). For ramos cell line experiments feature barcode 5’-TTGTCACGGTAATAA-3’ was used. For RM experiments feature barcodes 5’-TTGTCACGGTAATAA-3’ (AF647 bound probes), 5’ATCGCATTCTAAGAA3’ (AF488 bound probes), and 5’ATCTGCGCACATCTA3’ (AF546 negative bait) was used. Oligos were ordered from IDT and HPLC purified.

### Expression of recombinant streptavidin

One Shot^TM^ BL21(DE3) pLysE Chemically Competent E. coli (Invitrogen C656503) were transformed with plasmid of streptavidin modified with C-terminal cysteine with ampicillin resistance. Shortly, plasmid was added to a stock of bacteria. After 30 minutes on ice, bacteria were heat shocked for 30 seconds at 42° C. Afterward, bacteria were placed on ice for 5 minutes before culturing at 37° C for 60 min. A portion of the bacteria was plated onto an agar plate with 1:1000 ampicillin overnight at 37° C. A colony of bacteria was grown in lysogeny broth (LB) with 1:1000 ampicillin overnight at 37° C. Bacteria were diluted 1:1000 in Erlenmeyer flask and once an OD600 of 0.6 was reached, 0.4 mM IPTG was added for 4 hours before collecting bacteria at 4k x g for 15 min at 23° C.

To isolate and refold streptavidin, bacteria pellet was lysed with lysis buffer (30 mM Tris-HCl, 0.1% Triton X-100, 2 mM EDTA, pH 8) for 30 minutes on ice. Afterward, 12 mM MgSO4, 10 ug/ml DNAse 1, and 10 ug/ml RNAse A was added for 30 minutes on ice. Lysate was centrifuged at 14k x g for 20 minutes at 4° C, after which the pellet was washed three times with lysis buffer. Isolated streptavidin inclusion body was dissolved in denaturing buffer (6 M Guanidine Hydrochloride, pH 6.5). The dissolved streptavidin inclusion bodies were added to 3.5 kDa dialysis tubing and dialyzed overnight at 4° C in 6 M Guanidine Hydrochloride with 10 mM β-mercaptoethanol pH 1.5. Streptavidin monomers were refolded into tetramers by dialysis in refolding buffer (0.2 M sodium acetate, 10 mM β-mercaptoethanol, pH 6) over 8 hours at 4 C; this process was repeated 3 times with fresh refolding buffer. Refolded streptavidin was collected by 10 kDa size exclusion filtration.

### Conjugation of fluorophores to streptavidin

Recombinant streptavidin with C-terminal cysteine was buffer exchanged into cysteine buffer (100 mM Sodium Phosphate, 150 mM NaCl pH 7.2). Streptavidin was reduced with 50 molar excess tris(2-carboxyethyl)phosphine (TCEP) for 30 min at 23° C. Alexa fluorophores (AF) 488, 546, and 647 with maleimide group (Thermo Fisher) were dissolved in dimethyl sulfoxide (DMSO) and added at 10 molar excess to streptavidin for 2 hours at 23° C. Excess AF was removed through size exclusion spin columns (BIORAD 7326227) into spin buffer (150 mM NaCl, 100 mM Sodium Phosphate pH 6.5) according to manufacturer instructions. Concentration of streptavidin-AF was measured using Rapid Gold BCA Protein Assay (Thermo Fisher A55861) according to manufacture instructions.

### Conjugation of oligonucleotide barcodes to streptavidin

Streptavidin-AF was reduced with 50 molar excess TCEP for 30 min at 23° C. Afterwhich, 50 molar excess maleimide 6-hydrazinonicotinate acetone hydrazone (MHPH) (VectorLabs S-1009) dissolved in dimethylformamide (DMF) was added for 4 hours at 23° C. Excess MHPH was removed by 10 kD size exclusion filtration and buffer exchanged into conjugation buffer (150 mM Sodium Chloride, 50 mM Sodium Citrate pH 6). Additionally, oligonucleotide with 5’ amine modification was buffered exchanged into oligo buffer (100 mM Sodium Phosphate, 150 mM NaCl pH 8) and reacted to 25 molar excess sulfo succinimidyl 4-formylbenzoate (S-4FB) (VectorLabs S-1008) for 4 hours at 23° C. Excess S-4FB was removed by 3 kD size exclusion filtration and buffer exchanged into conjugation buffer. Streptavidin-AF and oligonucleotide were combined at 1 to 1 overnight at 23° C. Streptavidin-AF-oligonucleotide conjugate was purified from unconjugated streptavidin-AF and oligonucleotide using size exclusion chromatography Superdex 200 Increase 10/300 GL (Cytiva 28990944) on Akta pure chromatography system. Concentration of Streptavidin-AF-oligonucleotide conjugate was measured using Rapid Gold BCA Protein Assay (Thermo Fisher A55861) according to manufacturer instructions. Conjugate was visualized using NuPAGE Bis-Tris Mini Protein Gels, 4-12% (Thermo Fisher NP0321BOX) and stained with SYBR Gold Nucleic Acid Gel Stain (Thermo Fisher S11494) and Coomassie Brilliant Blue R-250 (Bio Rad 1610436) according to manufacturer instructions.

### Isolation of splenocytes from transgenic mouse spleen

Spleen was isolated from P14 transgenic mouse and placed in RPMI media (VWR 16750-070). Spleen was mashed using frosted glass slides and strained through 40 μm cell strainer. Red blood cells lysis (VWR 420301-BL) was added for 5 minutes at 4° C then diluted 5x with PBS. Splenocytes were suspended in RPMI media and strained through 40 μm cell strainer before use.

### Staining of splenocytes with pMHC tetramers

Recombinant Gp100-Db biotinylated monomers were tetramirized by adding streptavidin in 4 additions with 5 minute wait steps. Splenocytes were washed 3x with FACS buffer (1X PBS, 0.1% BSA, 2mM EDTA pH 7.4). Splenocytes were stained with Live/Dead aqua (Thermo Fisher L34957), PerCP/Cy5.5 anti-mouse CD8 (Biolegend Clone 53-6.7), and Gp100 tetramers (streptavidin-AF, streptavidin-AF-oligonucleotide, and Streptavidin-AF647 Thermo Fisher S21374) for 30 min at 4° C. Splenocytes were washed 3x with FACS buffer before running on Cytek Northern Light Flow Cytometer.

### Conjugation of streptavidin to biotinylated antigen

Biotinylated Env proteins and oligonucleotide bound streptavidin (DNA-STV) were centrifuged at 14,000 rcf for 10 min at 4C to spin down aggregates. Samples for conjugation were then pipetted from the top of each solution. 2.5ug of biotinylated Env (1ug/µL) were combined with 1ug of DNA-STV, and brought up to 10µL PBS (2:1 mass ratio of Env to DNA-STV), then mixed by pipetting up and down slowly 5x without introducing bubbles. Conjugation reactions were then incubated at 4 degrees for 1 hour away from light.

### Isolation of PBMCs

Peripheral blood lymphocytes were isolated from whole blood as described previously(57).

### Enrichment of antigen-specific B cells (FACS)

Up to 28 million PBMCs isolated from RM were sorted for 10x capture. Antigen specific memory B cells were classified through the following gates for sorting: lymphocytes and monocytes, singlet, live cells, CD3-CD14-CD16-, HLA-DR+, CD20+CD27+, IgM-IgG+, and BG505-STV-DNA-AF647+BG505-STV-DNA-AF488+ and Biotin-STV-DNA-AF546-. Cells were sorted using a BD FACSAria II instrument (BD Biosciences) at the Emory Vaccine Center Flow Cytometry Core at the ENPRC.

### Single-cell RNA-Sequencing

Single cell suspensions of FACS enriched memory B cells were prepared and loaded onto the 10X Genomics Chromium Controller using the Chromium NextGEM Single Cell 5’ Library & Gel Bead kit to capture individual cells and barcoded gel beads within droplets(58). For RM experiments, counting steps were skipped due to low number of antigen specific B cells isolated via FACS. VDJ and feature barcode libraries were prepared according to manufacturer instructions. They were then sequenced on an Illumina NovaSeq 6000 with a paired-end 26x91 configuration targeting a depth of 5,000 reads for both surface barcode libraries and VDJ libraires. Cell Ranger software was used to perform demultiplexing of cellular transcript data, as well as mapping and annotation of UMIs and transcripts for downstream data analysis.

### Single-cell RNA-Seq bioinformatic analysis of Memory B cells and Determination of LIBRA-seq Score

Cellranger v6.1.2 multi was used to obtain antigen barcode counts using the antigen barcode library and VDJ library. The VDJ reference that was used with multi was created using the fetch-imgt utility of cell ranger 6.0.2 for Macaca mulatta on 17th March 2023. The constant region sequences from Ramesh et al.(59) were added to the IMGT sequences (60). Since the multi option is only available with newer versions of cellranger which have specific requirements for VDJ reference, we used another VDJ reference comprised of heavy chain V gene sequences from Cirelli et al.(61), KimDB v1.1(62), IMPre(63) in addition to IMGT sequences to start with a comprehensive database for assembly with cellranger v3.1.0. The assembled sequences were once again annotated with the Cirelli et al database(61) using IgBLAST v1.21.0(64). The sequences were filtered to keep only those that were productive and with a predicted CDR3 region. After filtering Ig sequences, cells with only a single heavy and a single light chain were used downstream.

The antigen barcode data was processed using Seurat package v4.4.0(65). A seurat object was created after adding a pseudocount of 1 to the raw count data. The CLR normalization method was used with margin 2. Considering these were sorted to be antigen-specific cells, the thresholds for antigen barcodes were chosen empirically - 97th percentile for biotin and 3rd percentile for the two BG505-SOSIP barcodes. Using these thresholds, cells were classified as positive if the normalized values were higher for both BG505-SOSIP antigens and lower for biotin. Cells that were classified as double positive for BG505 and negative for biotin were considered as antigen-specific and subsequently used for downstream analysis. Clonal assignment was performed using a custom script with the family defined as cells having the same V genes, same J gene, same CDR3 length and CDR3 nucleotide identity >=70%. The V gene usage was calculated for clonal lineages using the countGenes function in the alakazam package(66).

The public clonotypes were defined using the same definition of clonal families as described above. The visualization for public clonotypes was generated using the Circos package(67).

To determine the neutralizing antibody lineage in the current dataset, the sequences for monoclonal antibodies of the neutralizing lineage(29) were downloaded from GenBank. These sequences were annotated using IgBLAST with the Cirelli et al database. The clonal lineage was determined using the definition described above.

For calculating SHM, we used the MUSA database (2025-02-05)(68). Only alleles that were found in both genomic and AIRR-Seq libraries of a given sample were used to create the IgBLAST v1.21.0 databases. The receptor_utils (http://pypi.org/project/receptor-utils/) package was used to create the J aux file. The MakeDB module from Immcantation v4.4.0(66) was used to create ChangeO tables from IgBLAST output. The observedMutations function was used from the shazam package(66) with the regionDefinition set to IMGT_V_BY_SEGMENTS and both frequency and combine set to TRUE to obtain SHM.

Cellranger v6.1.2 multi was used to obtain antigen barcode counts using the antigen barcode library and VDJ library. The reference that was used with multi was created using the fetch-imgt utility of cell ranger 6.0.2 for Macaca mulatta on 17th March 2023. The constant region sequences from Ramesh et al(59) were added to the IMGT sequences. Since the multi-option is only available with newer versions of Cellranger which have specific requirements for VDJ reference, we used another VDJ reference comprised of heavy chain V gene sequences from Cirelli et al.(61), KimDB v1.1(62), IMPre(63) in addition to IMGT sequences to start with a comprehensive database for assembly. To enable comparisons with the single-cell and bulk plasmablasts, the assembled sequences were once again annotated with the Cirelli et al.(61) database. The sequences were filtered to keep only productive sequences and cells with only a single heavy and a single light chain.

### LIBRA-Seq Scoring

Libra-Seq scores were determined as outlined by Setliff et al. previously(30). The antigen barcode data was processed using Seurat package. A Seurat object was created after adding 1 to the counts. Only cells that had a single productive heavy and light chain were retained. The CLR normalization method was used with margin 2. Considering these were sorted to be antigen-specific cells, the thresholds for antigen barcodes were chosen empirically - 97th percentile for biotin and 3rd percentile for the two BG505 SOSIP barcodes. Using these thresholds, cells were classified as positive if the normalized values were higher for both BG505 SOSIP antigens and lower for biotin. Cells that were classified as double positive for BG505 SOSIP and negative for biotin were subsequently used for downstream analysis.

IgBLAST v1.21.0 was used to annotate sequences and obtain AIRR-formatted outputs. The clones were defined as follows: (i) same V gene, (ii) same J gene, (iii) same CDR3 length and (iv) CDR3 nucleotide identity >= 70% for both heavy and light chains for determining clonal lineages.

### Monoclonal antibody generation

Variable domains were synthesized using Twist Bioscience. Next, they were directionally cloned into human mAb heavy chain (IgG1) and light chain (kappa/lambda) expression vectors (Genbank accession numbers FJ475055, FJ475056, and FJ517647). Following vector construction and sequence confirmation, heavy and light chain vectors were transiently co-transfected into Expi293F cells according to manufacturer’s instructions (Life Technologies). Antibodies were purified from cell culture supernatants using protein-A conjugated agarose beads (Pierce).

### TZM-bl neutralization assay

HIV-1 Env pseudoviruses were produced as previously described (46, 69, 70). Neutralization activity was assessed using the TZM-bl assay as previously described (46, 69, 70). In brief, 2000 infectious units of each Env pseudovirus were mixed with mAb at different concentrations and added to 96-well plates containing a TZM-bl monolayer. After 48 hr at 37 °C, cells were lysed, and luciferase activity was measured. Assays were performed in duplicate and independently repeated at least once. IC_50_ values were calculated in GraphPad Prism v10.

### BG505 Binding ELISA

MaxiSorp plates were coated with BG505[MOU1] .SOSIP.664 at 1µg/mL diluted in 50 mM carbonate buffer overnight at 4°C overnight at 4°C. The following day plates were washed with PBS Tween 0.05% then blocked with PBS 1% BSA for 90 minutes. Next, plates were incubated with mAbs diluted in PBS Tween 0.05% 1% BSA for 90 minutes. Plates were then washed and incubated with peroxidase conjugated goat anti human IgG (109-036-098) diluted in PBS Tween 0.5% 1% BSA for 90 minutes. Wells were developed with OPD substrate solution: 0.4 mg/ml of O-phenylenediamine (Sigma #P8787) dissolved into 50 mM citrate buffer (Sigma #P4560) with 30% H_2_O_2_. Plates were incubated with OPD substrate solution for 5 minutes. 100 ul of 1M HCl was added to stop the reaction and O.D. was recorded at 490 nm using the Bio-Rad IMark microplate reader.

### Data and Materials Availability

Data tables for expression counts for single-cell RNA-seq for memory B cells are deposited in NCBI’s Gene Expression Omnibus and are accessible through the Gene Expression Omnibus (GEO) under accession number (TBD). Custom scripts and supporting documentation on the RNA-seq analyses will be made available at https://github.com/BosingerLab/NHP_LIBRA-Seq. Reagents generated in this study may be requested from with a completed materials transfer agreement. All data needed to evaluate the conclusions in the paper are present in the paper or the Supplementary Materials.

## ACKNOWLEDGMENTS

We thank the Emory National Primate Research Center (ENPRC) Division of Animal Resources for providing support in animal care. We thank Kalpana Patel from the Environmental Health and Safety Office for BSL-3 training and safety oversight. We thank Dr. Daniel Lingwood and his group at the Ragon Institute for providing the engineered Ramos B cell lines. We thank Dr. Ivelin Georgiev and his group at Vanderbilt University for conceptualizing and developing the LIBRA-Seq approach and associated code.

## Author Contributions

CTE, GK, RA, and SEB conceptualized the study. AST produced and validated barcoded streptavidin probes with oversight from GK. A. Sahoo produced avi-tagged BG505.SOSIP.664.T332N trimers with oversight from RA. CTE conjugated barcoded streptavidin to biotinylated antigen. CTE and TT performed ramos cell assays and memory B cell staining. A. Saini at the Emory Primate/Vaccine/CFAR-Flow Cytometry Core performed sorting of antigen specific cells. AM and SAL performed 10x captures and library preparation. KP at the Emory Primate Center Genomics Core performed sequencing on 10x libraries. NR and AA performed bioinformatic scRNA-seq analyses. TCP and FM performed BG505 binding ELISAs with oversight from JW. CAD, EM and KC generated neutralization data. CTE analyzed flow cytometry, ELISA, and pseudovirus neutralization assay data and assembled figures. CTE, AST, and SEB wrote the manuscript with input from TT, and CAD reviewed and edited manuscript. Funding was acquired by SEB (U24AI120134), CAD (R01 AI174979), and RK (R01AI162267).

## LIST OF SUPPLEMENTARY MATERIALS

Fig. S1 to S7

Data Files S1-S4

